# Capsaicin inhibits arginine kinase and exerts anti-*Trypanosoma cruzi* activity

**DOI:** 10.1101/2020.06.23.167346

**Authors:** Edward A. Valera-Vera, Chantal Reigada, Melisa Sayé, Fabio A. Digirolamo, Mariana R. Miranda, Claudio A. Pereira

## Abstract

*Trypanosoma cruzi* is the causative agent of Chagas disease, considered within the list of twenty neglected diseases according to the World Health Organization. There are only two therapeutic drugs for Chagas disease, both of them unsuitable for the chronic phase, therefore the development of new drugs is a priority.

*T. cruzi* arginine kinase (TcAK) is a promising drug target since it is absent in humans and it is involved in cellular stress responses. In a previous study from our laboratory, possible TcAK inhibitors were identified through computer simulations, resulting in the best-scoring compounds cyanidin derivatives and capsaicin. Considering these results, in this work we evaluate the effect of capsaicin on TcAK activity and its trypanocidal effect. Although capsaicin produced a weak inhibition on the recombinant TcAK activity (IC_50_ ≈ 800 µM), it had a strong trypanocidal effect on epimastigotes and trypomastigotes (IC_50_ = 6.26 µM and 0.26 µM, respectively) being 20-fold more active on trypomastigotes than mammalian cells. Epimastigotes that overexpress TcAK were 37% more resistant to capsaicin than wild type parasites, suggesting that trypanocidal activity could be due, in part, to the enzyme inhibition. However, the difference between the concentrations at which the enzyme is inhibited and the parasite death is caused implies the presence of other targets. In this sense, the prohibitin-2 and calmodulin were identified as other possible capsaicin targets. Capsaicin is a strong and selective trypanocidal agent active in nanomolar concentrations, with an IC_50_ 57-fold lower than benznidazole, the drug currently used for treating Chagas disease.

## INTRODUCTION

Arginine kinase (AK, ATP:arginine phosphotransferase; EC 2.7.3.3) is an enzyme that catalyzes the reversible trans-phosphorylation between N-phospho-L-arginine and ATP [1]. Our research group identified an AK-mediated phosphagen system in the protozoan parasites *Trypanosoma cruzi* and *Trypanosoma brucei*, the etiological agents of Chagas disease and human sleeping sickness, respectively [2-4]. In *T. cruzi*, the AK pathway is involved in cellular response mechanisms to nutritional stress, radical oxygen species, and trypanocidal drugs [5-7]. The loss of flagellar AK in *T. brucei* reduces swim velocity and affects the infection in tsetse flies [8], and total elimination of the AK activity by RNA interference (RNAi) decreased parasite growth more than 90%, being lethal under oxidative stress conditions [9]. The presence of an arginine-based phosphagen and its biosynthetic pathway in these parasites, absent in the mammalian hosts, highlights the AK as a possible chemotherapy target against trypanosomiases such as Chagas disease (Pereira, 2014).

Chagas disease is a zoonosis that affects approximately 7 million people in Latin America, of which about 15%–30% develop the chronic clinical manifestations of the disease [10,11]. The epidemiological situation is aggravated if we consider the Chagas cases associated with migration towards non-endemic countries. Although early diagnosis and treatment are essential for a good prognosis, most of the infected people are detected late in the chronic phase. Currently there are only two available drugs for Chagas treatment; nifurtimox, introduced for clinical use in 1965, and benznidazole, introduced in 1971, both presenting severe side effects [12]. These trypanocidal treatments are less efficient in the chronic phase; for instance, benznidazole does not produce any improvement in the evolution of chronic chagas cardiomyopathy [13]. Although these drugs were discovered five decades ago, no new successful compounds have been developed to treat the disease, and those that reached clinical trials did not have the expected results [14].

The lack of new treatments is due in part to the little investment in research and development for this disease, raising the need for agile and low-cost strategies that allows the identification of new drugs and therapeutic alternatives. One of such strategies, the virtual screening, involves the use of computational tools to identify compounds with high chances of binding to a certain target [15]. In this sense, starting from the evidence of the green tea polyphenols catechin gallate and gallocatechin gallate inhibiting the *T. cruzi* AK (TcAK) [16,17], computational tools have been used to predict different plant-derived phenolic products as potential inhibitors of AK. Computer simulations validated by *in vitro* assays postulated rutin, a flavonoid glycoside, as a non-competitive inhibitor of the locust AK, binding to the enzyme mainly by hydrophobic interactions [18]. In a previous work from our research group, a virtual screening campaign performed on a database of phenolic compounds identified resveratrol as a TcAK inhibitor, which showed *in vitro* anti-parasite activity (IC_50_= 77 µM) and a moderate inhibition of the enzyme (IC_50_= 325 µM) [19]. In the same work, although not tested *in vitro*, the best-scored compounds from the virtual screening were three anthocyanidins, delphinidin, pelargonidin and cyanidin (polyphenolic plant pigments) and capsaicin [(E)-N-(4-hydroxy-3-methoxybenzyl)-8-methyl-6-nonenamide], a phenolic antioxidant compound produced by peppers.

Capsaicin has been tested for the treatment of cancer, pain, obesity, dermatitis, diabetes, and other conditions [20]. However, little is known about the effect of this compound on trypanosomatid parasites. Capsaicin is a specific inhibitor of the complex I of the respiratory chain in many organisms, and was used as a tool to study cellular respiration in *T. brucei* [21] and *Leishmania donovani* [22], however, in *T. cruzi* the role of mitochondrial complex I in NADH oxidation is very limited or null [23,24]. In addition, capsaicin was evaluated against *L. infantum* in combination with meglumine antimoniate, obtaining a moderate antiparasitic effect [25]. Considering the *in silico* predictions of capsaicin as a TcAK inhibitor, its use in the treatment of many human pathologies, and its activity in other trypanosomatids, in this work, we tested the effect of this compound on the TcAK activity and the parasite survival.

## MATERIALS AND METHODS

### Parasites and cells

*T. cruzi* epimastigotes of the Y strain were cultured at 28°C in BHT (brain-heart infusion-tryptose) medium supplemented with 10% fetal calf serum (FCS), 100 U/ml penicillin, 100 µg/ml streptomycin and 20 µg/ml hemin. Vero cells (African green monkey kidney) were cultured in MEM medium supplemented with 10% heat inactivated FCS, 0.15 % (w/v) NaHCO_3_, 100 U/ml penicillin and 100 U/ml streptomycin at 37°C in 5% CO_2_ atmosphere. Trypomastigotes were obtained from the extracellular medium of Vero infected cells as previously described [26].

### Arginine kinase overexpressing parasites

The full-length *T. cruzi* arginine kinase gene (GeneBank ID: AAC82390.1) was cloned into the pTREX expression plasmid and transfected into *T. cruzi* epimastigotes as previously described [5]. Transfected parasites were cultured in BHT medium containing 500 µg/ml of G418.

### Protein heterologous expression and purification

The full length TcAK gene was obtained by PCR amplification using genomic *T. cruzi* DNA as template and cloned into the pRSET-A expression vector (Invitrogen). Expression of recombinant AK was performed in *E. coli* strain BL21(DE3), cultivated in auto-induction medium (50 mM Na_2_HPO_4_, 50 mM KH_2_PO_4_, 10 mM NH_4_Cl, 5mM MgSO_4_, 4 mM NaCl, 0.2 mM CaCl_2_, 0.4% (v/v) glycerol, 0.05% (w/v) glucose, 0.05% (w/v) lactose, 1% (w/v) casein peptone, and 0.5% (w/v) yeast extract) supplemented with 0.2 mg/ml ampicillin. As the expressed protein formed inclusion bodies, bacteria were washed, lysed by sonication, and the insoluble phase of the lysate was kept and washed three times with 50 mM Tris–HCl buffer pH 8.0, denatured with 6 M urea, and refolded by dilution 1:20 in 50 mM Tris–HCl buffer pH 8.0, 10% (v/v) glycerol, 5% (w/v) sucrose during 72 h at 4°C. Purity of the protein was checked by SDS-PAGE followed by Coomassie blue staining.

### Arginine kinase activity assays

TcAK activity was measured following the ADP production with coupled enzyme reactions as previously described [27]. Briefly, an 8 μg aliquot of recombinant TcAK was added to 100 μl of the reaction mixture (100 mM Tris–HCl buffer pH 8.2, 1.5 mM ATP, 1.5 mM phosphoenolpyruvate, 1.5 mM MgCl_2_, 0.5 mM DTT, 0.3 mM NADH, 10 mM L-arginine, 5 U/ml pyruvate kinase and 5 U/ml lactate dehydrogenase) in a 96-well plates. The enzyme activity was determined measuring the decrease in absorbance at 340 nm due to the oxidation of NADH.

### Trypanocidal activity assays

Activity on epimastigotes was tested using 10^7^ cells/ml epimastigote cultures in 24-well plates treated with 0–50 µM of capsaicin during 72 h. Activity against trypomastigotes was evaluated using 10^6^ cells/ml in 96-well plates incubated at 37°C for 24 h in the presence of 0–2.5 µM of capsaicin. Cellular density was determined by cell counting in a Neubauer chamber.

### Vero cell viability assay

Cytotoxicity against Vero cells was assayed by the crystal violet staining assay. The cells (10^4^ cells well) were incubated at 37 for 24 h in 96-well plates with 0–400 µM of capsaicin. Then, cells were fixed for 15 min in methanol, and stained with 0.5 % (w/v) crystal violet. After washing with water and drying, the crystal violet staining the cells was solubilized with 200 μl of methanol, and the absorbance was measured at 570 nm.

### Statistics and data analysis

IC_50_ values were obtained by non-linear regression of dose-response logistic functions, using GraphPad Prism 6.01. All experiments were performed in triplicate and results are presented as mean ± standard deviation (SD).

### Alternative targets search

Other possible molecular targets of capsaicin were searched using the following bioinformatics tools and databases: PASS (http://www.akosgmbh.de/pass/) [28], Similarity ensemble approach (SEA, http://sea.bkslab.org/) [29], Swiss Target Prediction (http://www.swisstargetprediction.ch/) [30], Target Hunter (https://www.cbligand.org/TargetHunter/) [31], KEGG Compounds database (https://www.genome.jp/kegg/compound) [32], PubChem (https://pubchem.ncbi.nlm.nih.gov/) and DrugBank (https://www.drugbank.ca/).

## RESULTS

### Inhibition of TcAK

In order to determine if capsaicin can act as an TcAK inhibitor, enzyme activity assays were performed using a concentration range between 0 – 2 mM capsaicin in the presence of saturating concentrations of the substrates L-arginine and ATP. As Fig. 1A shows, TcAK activity in presence of capsaicin decreased in a dose-dependent manner, reaching 50% inhibition only at high concentrations of capsaicin of approximately 0.8 mM.

**Figure 1.**
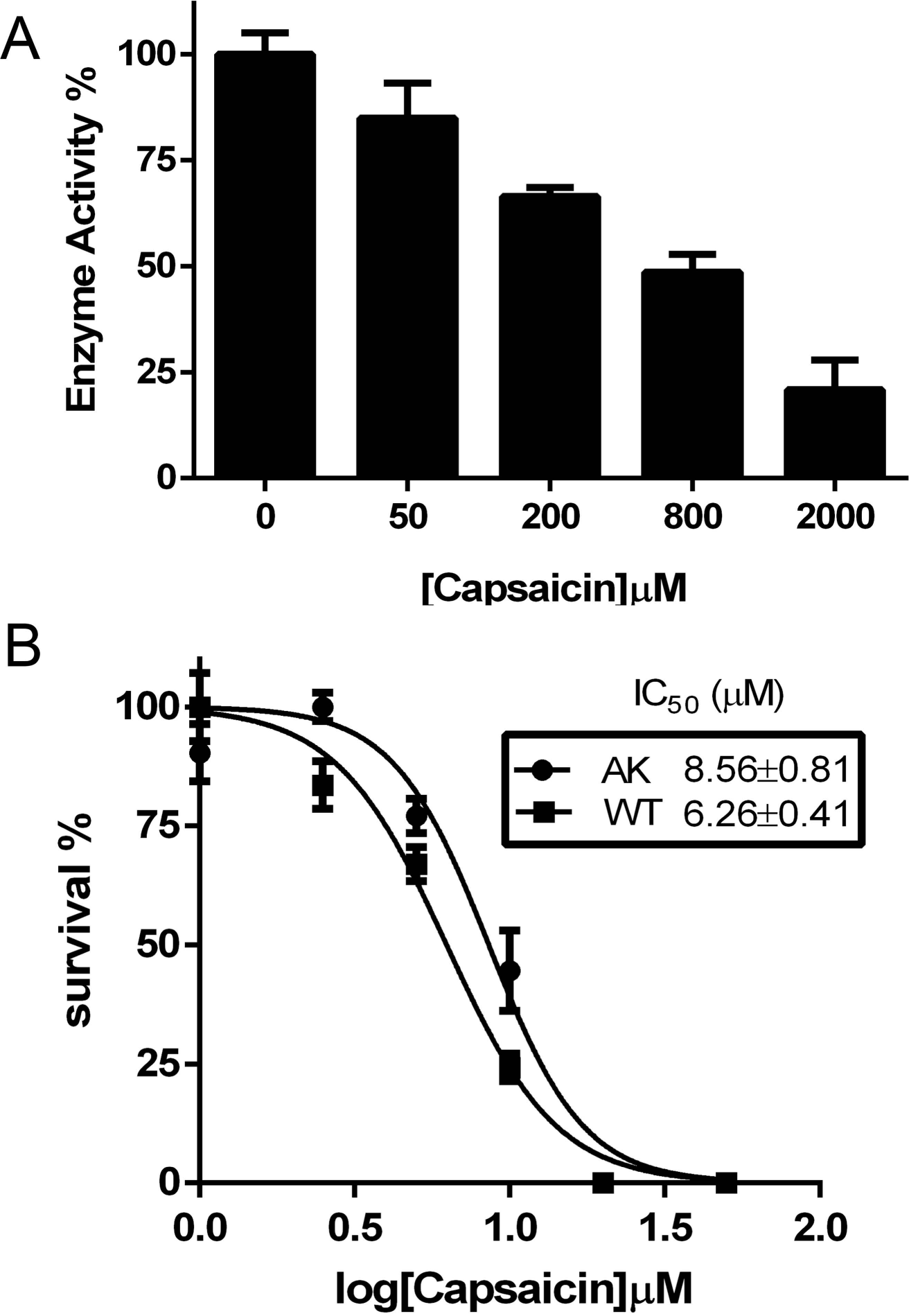
Effect of capsaicin on TcAK activity and epimastigotes viability. A) TcAK activity was measured by coupled enzymes method in presence of 0 - 2 mM capsaicin, under saturating concentration of arginine and ATP, using a recombinant purified enzyme as described under “Materials and Methods”. Activity was expressed as percentage of the control without capsaicin. B) Trypanocidal effect of capsaicin treatment using a concentration range of 0 - 50 µM during 72 h, in wild type (■) and TcAK overexpressing (●) epimastigotes. I _50_s were obtained by non-linear regression, expressed as the mean ± standard deviation and corresponds to three independent experiments.

### Trypanocidal effect on epimastigotes

To evaluate the effect of capsaicin on the replication of the insect-infecting stage of *T. cruzi*, epimastigote cells were exposed to the compound during 72 h in a concentration range of 0 – 50 µM. Results showed that capsaicin inhibited the epimastigotes growth with an IC_50_ (concentration of compound which gave 50% relative number of parasites compared to the untreated control) of 6.26 (± 0.41) µM (Fig. 1B).

Considering that AK is an important component of stress responses in *T. cruzi*, we evaluated the role of this enzyme in the mechanism of action of the capsaicin-mediated trypanocidal effect. Thus, we compared the IC_50_ calculated for wild type epimastigotes and transgenic parasites overexpressing AK. Remarkably, epimastigotes overexpressing AK were 37% more resistant to capsaicin than control parasites, with an IC_50_ of 8.56 (± 0.81) µM (Fig. 1B).

As the concentrations of capsaicin inhibiting the enzyme are two orders of magnitude higher than the concentrations killing the parasite, the results suggest that the effect on epimastigote viability is not a direct consequence of TcAK inhibition, but the higher resistance of the overexpressing cells indicates that this pathway could be involved in the response to the capsaicin-mediated trypanocidal effect.

### Trypanocidal effect on trypomastigotes

The trypanocidal effect of capsaicin was tested in bloodstream trypomastigotes, a therapeutically relevant stage of the *T. cruzi* life cycle. Interestingly, when trypomastigotes were exposed to capsaicin concentrations up to 2.5 µM, the calculated IC_50_ was 0.26 (± 0.02) µM demonstrating that trypomastigotes are more sensitive to capsaicin than epimastigotes (Fig. 2A). Moreover, the compound presented an IC_50_ 57-fold lower than the drug benznidazole, being 14.9 (± 1.5) µM when tested in similar conditions [33]. Finally, the cytotoxicity of capsaicin was assayed using the mammalian Vero cell line (Fig. 2B) and the calculated IC_50_ was 5.26 (± 0.44) µM, representing a selectivity index (IC_50_ Vero / IC_50_ trypomastigotes) of 20.23.

**Figure 2.**
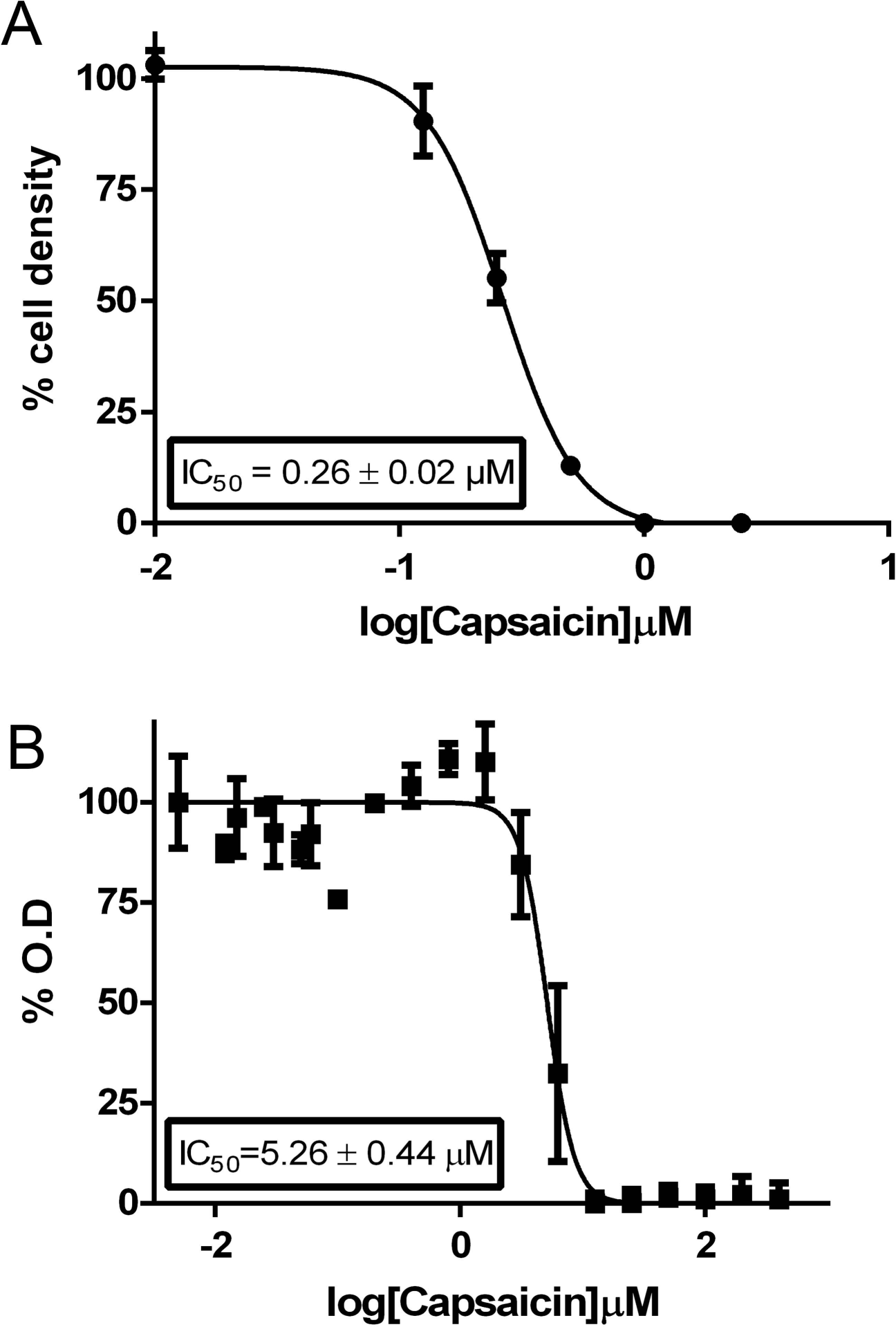
Trypanocidal activity of capsaicin on trypomastigotes. Dose-response curves of capsaicin over trypomastigotes of *T. cruzi* (A) and Vero cells (B). Treatments were performed during 24 h as indicated under “Materials and Methods”. % OD is the percentage of absorbance at λ = 570 nm of the crystal violet (vital staining) respect to the control without treatment. IC_50_s were obtained by non-linear regression, expressed as the mean ± standard deviation and corresponds to three independent experiments.

### Prediction of other capsaicin targets in *T. cruzi*

As previously mentioned, considering the differences between the IC_50_ obtained for AK inhibition and the trypanocidal activity on epimastigotes and trypomastigotes (about 100 and 3,000-fold lower, respectively) it can be hypothesized that other targets of capsaicin, besides TcAK, are present in *T. cruzi*. Thus, applying seven *in silico* “target fishing” tools and databases two protein candidates, that could interact with capsaicin, were obtained. A search in the DrugBank database provided information about the binding of capsaicin to the human prohibitin-2 (PHB2), a protein involved in the maintenance of mitochondrial morphology and the control of apoptosis. This interaction induces PHB2 translocation from the inner mitochondrial membrane, where it is normally localized, to the nucleus, triggering apoptosis at multiple levels [34]. Regarding trypanosomatids, in *T. brucei* PHB1 and PHB2 are located in the mitochondrion and seem to be essential for mitochondrial translation, to maintain the flagellar morphology and are also involved in apoptosis [35,36]. It was postulated in *L. major* that PHB1 and PHB2 protect membranes and DNA against superoxide ions in oxidative stress conditions, increasing the survival capacity of the parasites [37].

A search in the PubChem database showed that capsaicin binds to human calmodulin at nanomolar concentrations, competitively inhibiting its interaction with TRPV1, a ionotropic receptor involved in pain stimulus [38]. Interestingly, there is a report of another flavonoid (5-methoxy-6,7-methylenedioxyflavone) affecting the Ca^2+^ regulation in trypanosomatids [39]. Other bioinformatics tools used (see “Materials and Methods”) did not give any hits or they were not applicable to trypanosomatids.

All this evidence highlights PHB2 and calmodulin as possible targets for capsaicin in *T. cruzi*.

## DISCUSSION

Phenolic compounds derived from natural products have been intensively studied to treat a wide variety of conditions, including cancer, inflammation, pain, ageing, cardiomyopathy, hepatotoxicity, obesity, viral and bacterial infections, dermatitis, among other pathologies [40].

The enzyme arginine kinase, completely absent in mammals, is involved in stress responses through the maintenance of the cell energy homeostasis. The essentiality of this enzyme was demonstrated by RNAi in *T. brucei* and, by different pharmacological approaches, in *T. cruzi* [9,17]. There are two identified chemical groups of TcAK inhibitors: arginine structural analogs and plant-produced phenolic compounds [19,27]. In a previous work we performed a virtual screening campaign searching potential *T. cruzi* AK inhibitors in a database of antioxidant phenolic compounds, where resveratrol was selected for further characterization [19]. Although capsaicin scored higher than resveratrol as a possible TcAK inhibitor in the virtual screening (−8.56 and −6.96 kcal/mol, respectively), the experimental validation showed that capsaicin achieved 50% enzyme inhibition at concentrations 2.5-fold higher than the reported for resveratrol. Regarding the trypanocidal activity, capsaicin performed better than resveratrol, with potency 15-fold higher in epimastigotes and 300-fold in trypomastigotes. According to the results herein obtained, capsaicin acts as a moderate inhibitor of *T. cruzi* AK. However, the trypanocidal potency of this compound is 2 to 3 orders of magnitude higher than the TcAK inhibition, suggesting that capsaicin has other targets besides TcAK. In addition, TcAK overexpression increases IC_50_ of capsaicin suggesting that although it is not the main target, it is probably involved in resistance mechanisms to capsaicin-mediated cytotoxicity.

Searching in different compound databases, two probable targets were identified, the protein PHB2, a multifunctional protein involved in apoptosis and oxidative stress protection, and calmodulin, a calcium-binding messenger protein expressed in all eukaryotic cells. The binding of capsaicin to human PHB2 and calmodulin were previously demonstrated [34,38] and both proteins are present in trypanosomatid organisms. Although other targets have not been determined with certainty, there is convincing evidence of the multitarget nature of capsaicin trypanocidal activity.

Capsaicin can be considered a potent and selective anti-*T. cruzi* agent since it is active in nanomolar concentrations, 57-fold more potent than benznidazole [41], the drug currently used to treat Chagas disease. In addition, orally administered free capsaicin suspension and capsaicin loaded liposomes in rats at a single dose of 90 mg.kg^-1^ reach a plasma concentration of about 2.6 µM that is progressively reduced to 50% at 2 h or 24 h for free or liposome formulation of capsaicin, respectively [42]. These data are particularly relevant since the calculated IC_50_ for *T. cruzi* trypomastigotes is 10-fold lower than the maximum concentration achieved in plasma, suggesting that capsaicin could be a good candidate for an oral treatment of trypanosomiasis.

Finally, other advantages of capsaicin to be used as a trypanocidal drug include that can be administered orally (free or in liposomes), it is very stable at room temperature, easily available at a low market price (about about USD 270 per kg) and its pharmacological properties are widely known.

## Acknowledgements

This work was supported by Consejo Nacional de Investigaciones Científicas y Técnicas, Agencia Nacional de Promoción Científica y Tecnológica (FONCyT PICT 2015–0539, 2018-1801 and 2018-1871). The research leading to these results has, in part, received funding from UK Research and Innovation via the Global hallenges Research Fund under grant agreement ‘A Global Network for Neglected Tropical Diseases’ grant number MR/P027989/1. AP and MRM are members of the career of the scientific investigator; EVV and CR are research fellows from CONICET; FAD is a scientific technician from CONICET, and MS is PDRA from the A Global Network for Neglected Tropical Diseases.

## Notes

### Competing Interest Statement

The authors have declared no competing interest.

